# PSI controls *tim* splicing and circadian period in *Drosophila*

**DOI:** 10.1101/504282

**Authors:** Lauren Foley, Jinli Ling, Radhika Joshi, Naveh Evantal, Sebastian Kadener, Patrick Emery

## Abstract

The *Drosophila* circadian pacemaker consists of transcriptional feedback loops subjected to both post-transcriptional and post-translational regulation. While post-translational regulatory mechanisms have been studied in detail, much less is known about circadian post-transcriptional control. To have a better understanding of the role and mechanisms of circadian post-transcriptional regulation, we targeted 364 RNA binding and RNA associated proteins with RNA interference. Among the 43 genes we identified was the alternative splicing regulator P-element somatic inhibitor (PSI). PSI downregulation shortens the period of circadian rhythms both in the brain and in peripheral tissues. Interestingly, we found that PSI regulates the thermosensitive alternative splicing of *timeless (tim)*, promoting splicing events favored at warm temperature over those increased at cold temperature. Moreover, the period of circadian behavior was insensitive to PSI downregulation when flies could produce functional TIM proteins only from a transgene that cannot form the thermosensitive splicing isoforms. Therefore, we conclude that PSI regulates the period of *Drosophila* circadian rhythms through its modulation of the *tim* splicing pattern.

## Introduction

Circadian rhythms are the organism’s physiological and behavioral strategies for coping with daily oscillations in environment conditions. Inputs such as light and temperature feed into a molecular clock via anatomical and molecular input pathways and reset it every day. Light is the dominant cue for entraining the molecular clock, but temperature is also a pervasive resetting signal in natural environments. Paradoxically, clocks must be semi-resistant to temperature: they should not hasten in warm summer months or lag in the winter cold (this is called temperature compensation), but they can synchronize to the daily rise and fall of temperature (temperature entrainment) (Pittendrigh, 1960). Not only can temperature entrain the clock, it also has a role in seasonal adaptation by affecting the phase of behavior (see for example Majercak et al., 1999).

Molecular circadian clocks in eukaryotes are made up of negative transcriptional feedback loops (Dunlap, 1999). In *Drosophila*, the transcription factors CLOCK (CLK) and CYCLE (CYC) bind to E-boxes in the promoters of the clock genes *period (per)* and *timeless (tim)* and activate their transcription. PER and TIM protein accumulates in the cytoplasm where they heterodimerize and enter the nucleus to feedback and repress the activity of CLK and CYC and thus down-regulate their own transcription (Hardin, 2011). This main loop is strengthened by a scaffolding of interlocked feedback loops involving the transcription factors *vrille* (*vri*), *PAR domain protein 1* (*Pdp1*) and *clockwork orange* (*cwo*). Post-translational modifications are well-established mechanisms for adjusting the speed and timing of the clock (Tataroglu and Emery, 2015).

Increasing evidence indicates that post-transcriptional mechanisms controlling gene expression are also critical for the proper function of circadian clocks in many organisms. In *Drosophila*, the post-transcriptional regulation of *per* mRNA has been best studied. *per* mRNA stability changes as a function of time (So and Rosbash, 1997). In addition, *per* contains an intron in its 3’UTR (dmpi8) that is alternatively spliced depending on temperature and lighting conditions (Majercak et al., 1999; Majercak et al., 2004). On cold days, the spliced variant is favored, causing an advance in the accumulation of *per* transcript levels as well as an advance of the evening activity peak. This behavioral shift means that the fly is more active during the day when the temperature would be most tolerable in their natural environment. The temperature sensitivity of dmpi8 is due to the presence of weak non-canonical splice sites. However, the sensitivity of the underlying baseline splicing is affected by 4 single nucleotide polymorphisms (SNPs) in the *per* 3’UTR that vary in natural populations and form two distinct haplotypes (Low et al., 2012; Cao and Edery, 2017). Also, two *Drosophila* species that remained in Africa lack temperature sensitivity of dmpi8, while two species that followed human migration do (Low et al., 2008). Furthermore, Zhang *et al* recently demonstrated that the increase in dmpi8 splicing efficiency caused by one of the two haplotypes is mediated by the *trans*-acting splicing factor B52 (Zhang et al., 2018). *per* is also regulated post-transcriptionally by the TWENTYFOUR-ATAXIN 2 translational activation complex (Lim et al., 2011; Lim and Allada, 2013a; Zhang et al., 2013; Lee et al., 2017). This complex works by binding to *per* mRNA as well as the cap-binding complex and poly-A binding protein. This may enable more efficient translation by promoting circularization of the transcript. Interestingly, this mechanism appears to be required only in the circadian pacemaker neurons. Non-canonical translation initiation has also been implicated in the control of PER translation (Bradley et al., 2012). Regulation of PER protein translation has also been studied in mammals, with the mammalian LARK homolog (RBM4) being a critical regulator of mPER1 expression (Kojima et al., 2007). In flies however, LARK regulates the translation of DBT, a PER kinase (Huang et al., 2014). miRNAs have emerged as important critical regulators of circadian rhythms in *Drosophila* and mammals, affecting the circadian pacemaker itself, as well as input and output pathways controlling rhythmic behavioral and physiological processes (Lim and Allada, 2013b; Tataroglu and Emery, 2015)

RNA associated proteins (RAPs) include proteins that either bind directly or indirectly to RNAs. They mediate post-transcriptional regulation at every level. Many of these regulated events – including alternative splicing, splicing efficiency, mRNA stability, and translation – have been shown to function in molecular clocks. Thus, to obtain a broad view of the *Drosophila* circadian RAP landscape and its mechanism of action, we performed an RNAi screen targeting 364 of these proteins. This led us to discover a role for the splicing factor *P-element somatic inhibitor (Psi)* in regulating the pace of the molecular clock through alternative splicing of *tim*.

## Results

### An RNAi screen for RNA associated proteins controlling circadian behavioral rhythms

Under constant darkness conditions (DD) flies have an intrinsic period length of about 24 hours. To identify novel genes that act at the post-transcriptional level to regulate circadian locomotor behavior, we screened 364 genes that were annotated in either Flybase or the RNA Binding Protein Database as RNA binding or involved in RNA associated processes using period length as a readout of clock function (Dataset S1). We avoided many, but not all, genes with broad effects on gene expression, such as those encoding essential splicing or translation factors. When possible, we used at least two non-overlapping RNAi lines from the TRiP and VDRC collections. RNAi lines were crossed to two different GAL4 drivers: *tim-GAL4* (Kaneko et al., 2000) and *Pdf-GAL4* (Renn et al., 1999) each combined with a *UAS-dicer-2* transgene to enhance the strength of the knockdown (Dietzl et al., 2007). These combinations will be abbreviated as *TD2* and *PD2*, respectively. *tim-GAL4* drives expression in all cells with circadian rhythms in the brain and body (Kaneko et al., 2000), while *Pdf-GAL4* drives expression in a small subset of clock neurons in the brain: the PDF-positive small (s) and large (l) LNvs (Renn et al., 1999). Among them, the sLNvs are critical pacemaker neurons that drive circadian behavior in DD (Renn et al., 1999; Stoleru et al., 2005). In the initial round of screening, we tested the behavior of 4-8 males for each RNAi line crossed to both *TD2* and *PD2* (occasionally, fewer males were tested if a cross produced little progeny). We also crossed some RNAi lines to *w^1118^* (+) flies (most were lines selected for retest, see below). We noticed that RNAi/+ control flies for the TRiP collection were 0.3 hr shorter than those of the VDRC collection (Figure 1A). Furthermore, the mean period from all RNAi lines crossed to either *PD2* or *TD2* was significantly shorter for the TRiP collection than for the VDRC collection (Figure 1A) (0.2 hr, *TD2* crosses; 0.5 hr, *PD2* crosses). We also found that many of the VDRC KK lines that resulted in long period phenotypes when crossed to both drivers, (but especially when crossed to *PD2)*, contained insertions in the 40D locus (VDRC annotation). It has been shown that this landing site is in the 5’UTR of *Tiptop (tio)* and can lead to non-specific effects in combination with some GAL4 drivers, likely due to misexpression of *tio* (Green et al., 2014; Vissers et al., 2016). Indeed, when we crossed a control line that contains a UAS insertion at 40D (*40D-UAS*) to *PD2*, the progeny also had a ca. 0.6hr longer period relative to the *PD2* control (Figure 1B). Thus, in order to determine a cutoff for candidates to be followed-up, we analyzed the data obtained in our screen from the TRiP, VDRC, and the 40D KK VDRC lines independently (Figure 1C). These data are represented in two overlaid histograms that show period distributions: one for the *TD2* crosses and one for the *PD2* crosses. We chose a cutoff of two standard deviations (SD) from the mean period length for each RNAi line set. The lines that showed circadian behavior with periods falling outside the two standard deviation cutoff were selected for repeat. We also chose to repeat a subset of lines below the cutoff that were of interest and showed period lengthening or shortening, as well as lines that were highly arrhythmic in constant darkness (DD) or had an abnormal pattern of behavior in a light-dark cycle (LD). After a total of three independent experiments, we ended up with 43 candidates (Table 1) that passed the period length cut offs determined by the initial screen; 31 showed a long period phenotype, while 12 had a short period. One line showed a short period phenotype with *PD2* but was long with *TD2* (although just below the 2-SD cutoff). Although loss of rhythmicity was also observed in many lines (Dataset S1), we decided to focus the present screen on period alterations to increase the probability of identifying proteins that regulate the circadian molecular pacemaker. Indeed, a change in the period length of circadian behavior is most likely caused by a defect in the molecular pacemaker of circadian neurons, while an increase in arrhythmicity can also originate from disruption of output pathways, abnormal development of the neuronal circuits underlying circadian behavioral rhythms, or cell death in the circadian neural network, for example.

**Table 1:**
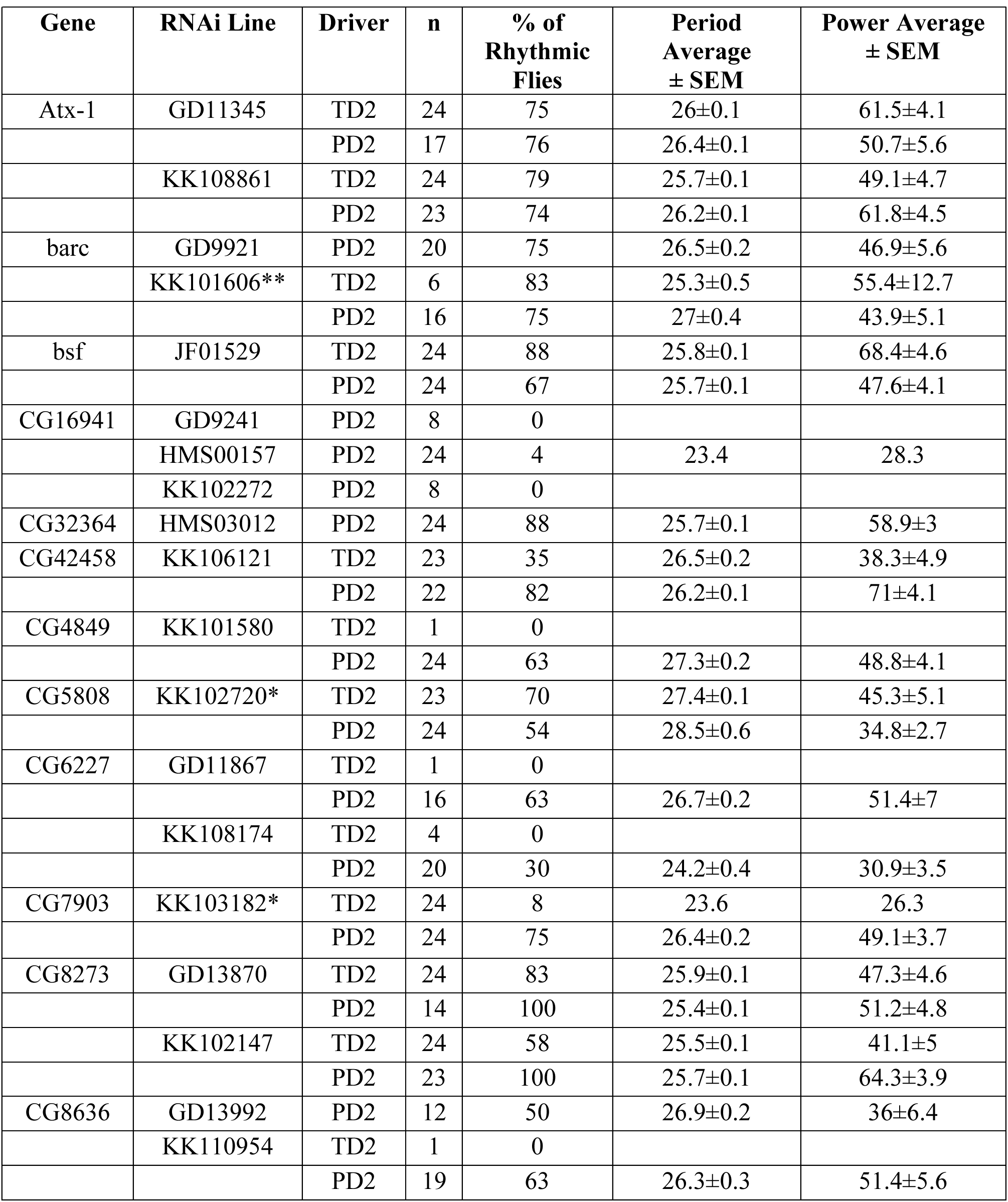
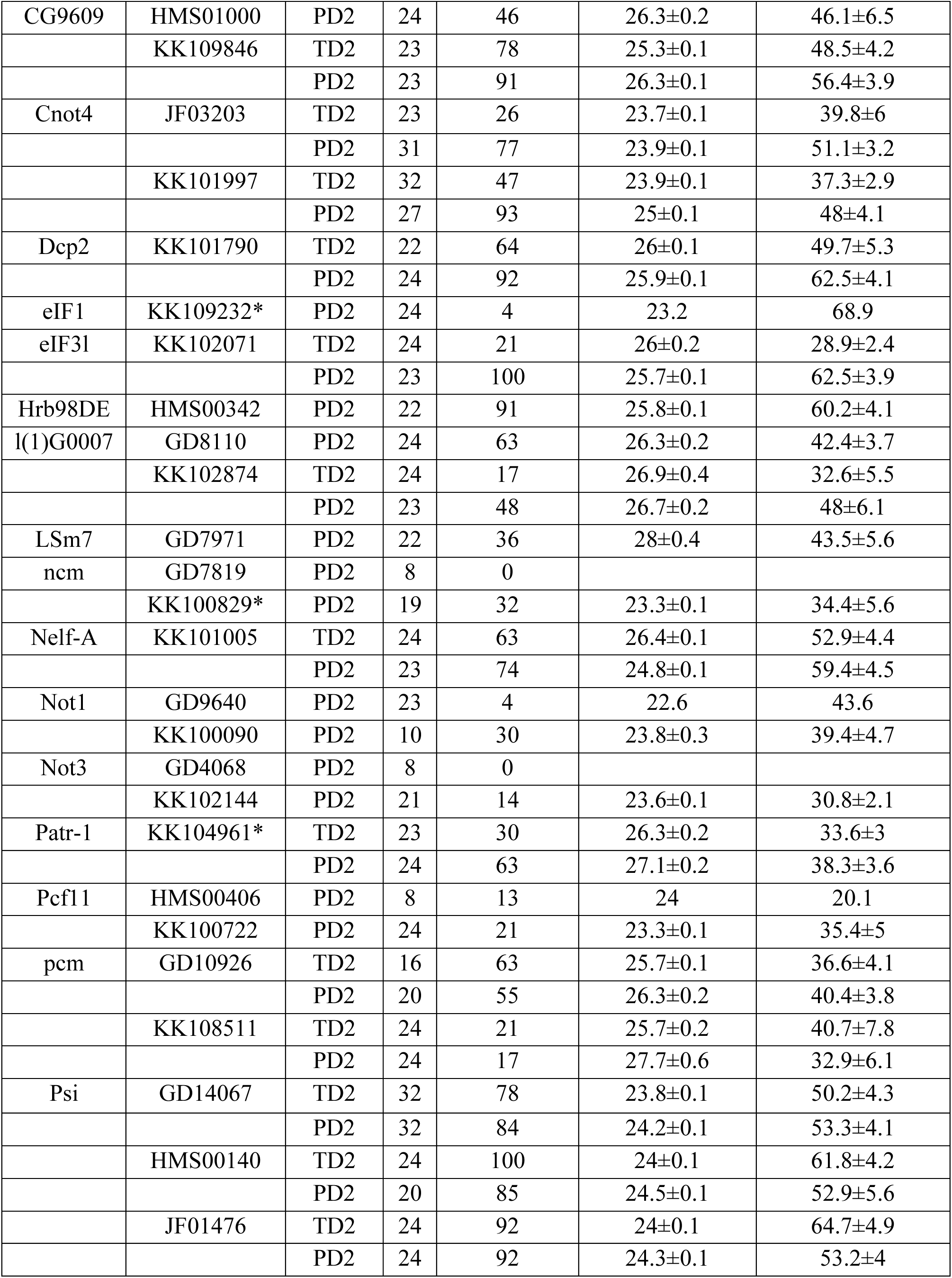
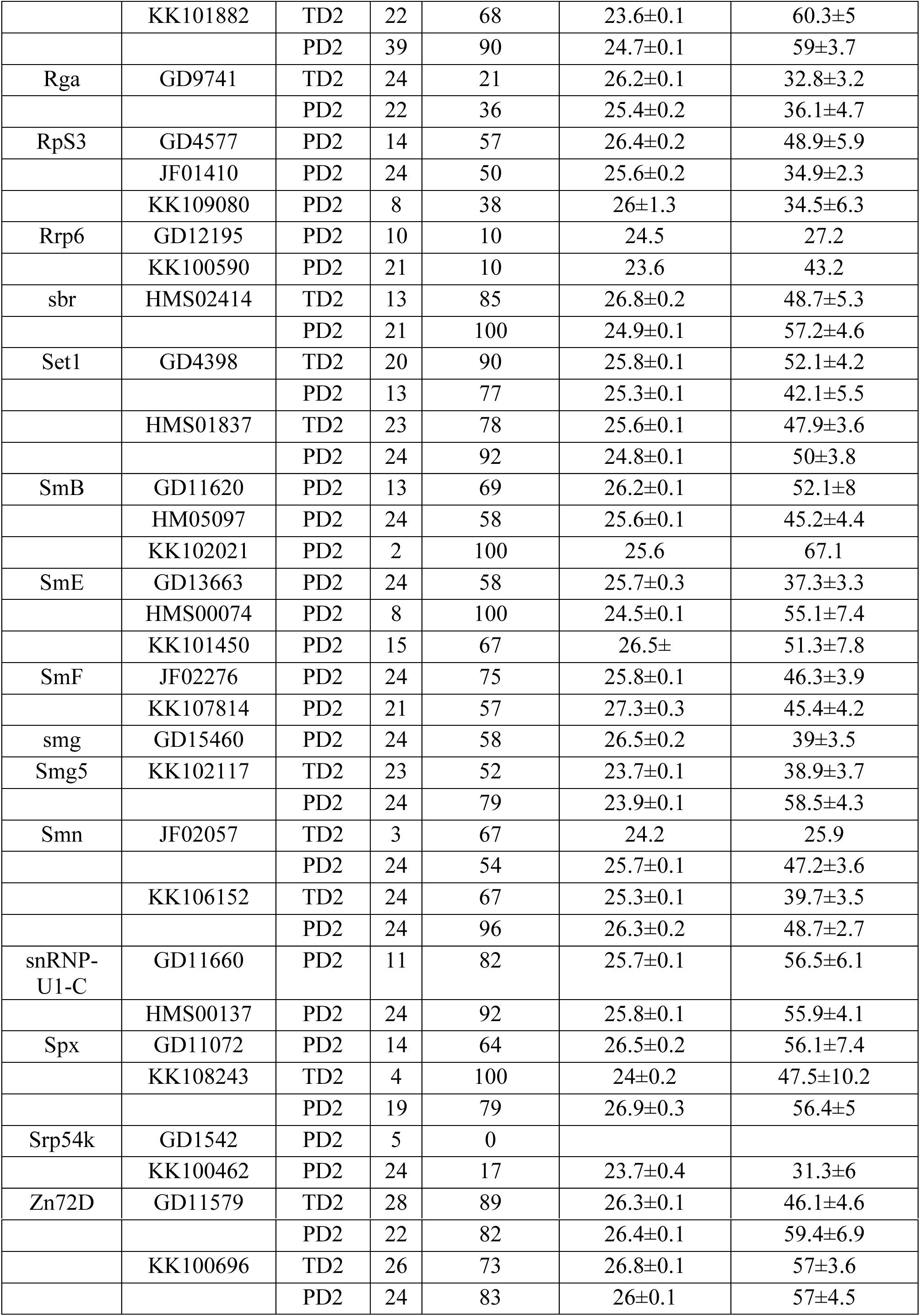
Circadian behavior in DD of screen candidates

**\* Line contains insertion at 40D. ** Unknown if line contains insertion at 40D.**

**Figure 1.**
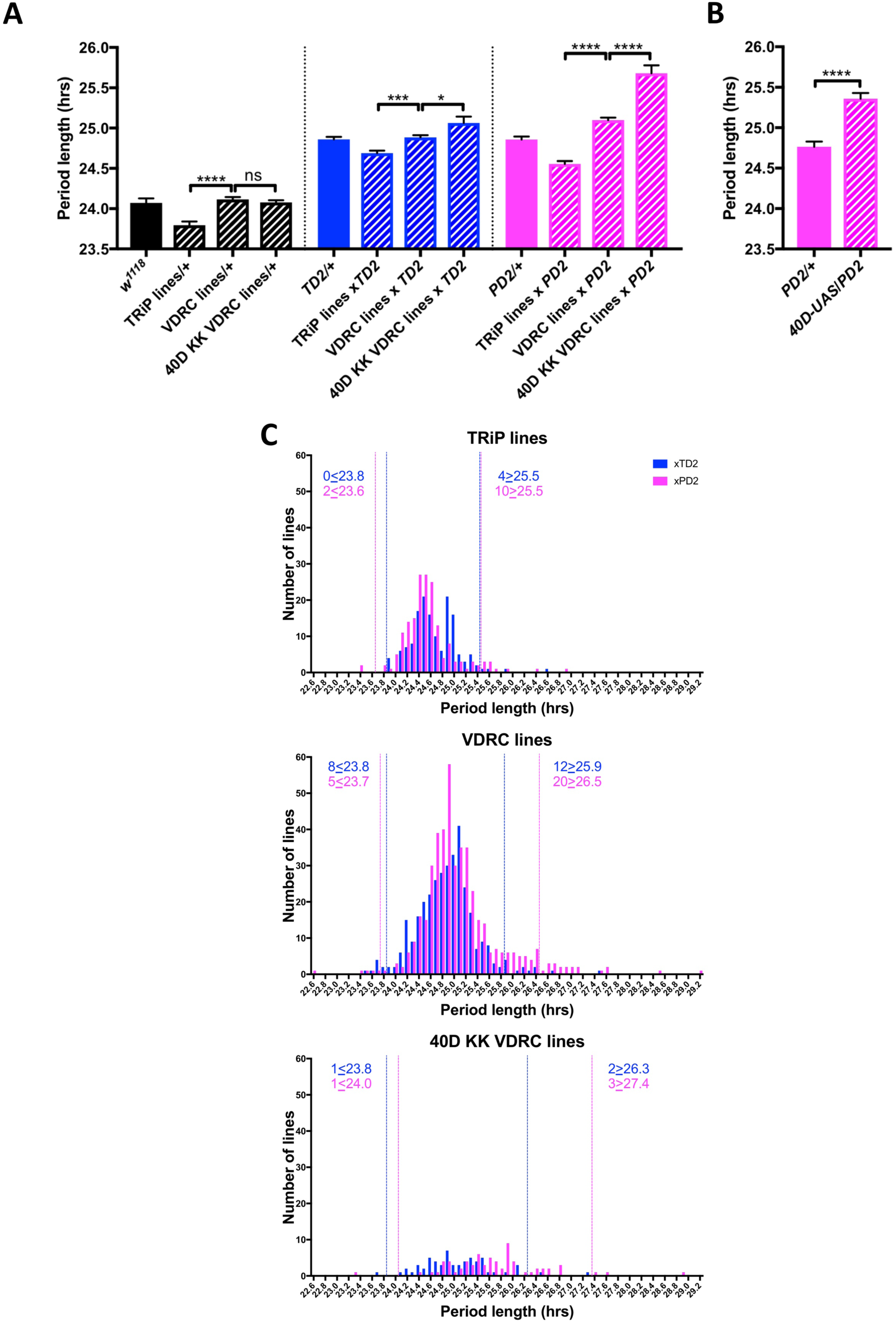
An RNAi screen of RNA associated proteins identifies long and short period hits. (A-B) Background effect of TRiP and VDRC collections on circadian period length. Circadian period length (hrs) is plotted on the y axis. RNAi collection and genotypes are labeled. Error bars represent SEM. (A) Left group (black bars): Patterned bars are the average of period lengths of a subset of RNAi lines in the screen crossed to *w^1118^* (*TRiP/+* N=17 crosses*, VDRC/+* N=46 crosses, *40D KK VDRC/+* N=20 crosses). Solid bar is the *w^1118^* control (N=20 crosses). Middle group (blue bars): Patterned bars are the average of period lengths of all RNAi lines in the screen crossed to *tim-GAL4, UAS-Dicer2 (TD2)* (*TRiP/TD2* N=151 crosses*, VDRC/TD2* N=340 crosses, *40D KK VDRC/TD2* N=61 crosses). Solid bar is the *TD2/+* control (N=35 crosses). Right group (magenta bars): Patterned bars are the average of period lengths of all RNAi lines in the screen crossed to *Pdf-GAL4, UAS-Dicer2 (PD2)* (*TRiP/PD2* N=176 crosses*, VDRC/PD2* N=448 crosses, *40D KK VDRC/PD2* N=69 crosses). Solid bar is the *PD2/+* control (N=36 crosses). One-way analysis of variance (ANOVA) followed by Tukey’s multiple comparison test: *p<0.05, ***p<0.001, ****p<0.0001. Note that the overall period lengthening, relative to wild-type (*w^1118^*), when RNAi lines are crossed to *TD2* or *PD2* is most likely a background effect of our drivers, while the period differences between the TRiP (shorter) and VDRC (longer) collections is most likely a background effect of the RNAi lines themselves. There is also a lengthening effect of the 40D insertion site in the VDRC KK collection that cannot be explained by a background effect, as it is not present in the RNAi controls (Left panel). Instead the lengthening was only observed when these lines were crossed to our drivers. A modest effect was seen with *TD2* (middle panel) and a larger effect was seen with *PD2* (right panel). (B) The period lengthening effect of the VDRC 40D KK lines is likely due to overexpression of *tio*, as we observed lengthening when a control line that lacks a RNAi transgene, but still has a UAS insertion in the 40D (*40D-UAS*) locus was crossed to *PD2.* N=32 flies per genotype, ****p<0.0001, Student’s t-test. (C) Histogram of period lengths obtained in the initial round of screening. Number of lines per bin is on the y axis. Binned period length (hrs) is on the x axis. Bin size is 0.1 hrs. *TD2* crosses are in blue and *PD2* crosses are in magenta. Dashed lines indicate our cutoff of 2 standard deviations from the mean. Number of crosses that fell above or below the cutoff is indicated. Top panel: TRiP lines. 0 lines crossed to *TD2* and 2 lines crossed to *PD2* gave rise to short periods and were selected for repeats. 4 lines crossed to *TD2* and 10 lines crossed to *PD2* gave rise to long periods and were selected for repeats. Middle panel: VDRC lines. 8 lines crossed to *TD2* and 5 lines crossed to *PD2* gave rise to short periods and were selected for repeats. 12 lines crossed to *TD2* and 20 lines crossed to *PD2* gave rise to long periods and were selected for repeats. Bottom panel: VDRC 40D KK lines. 1 line crossed to *TD2* and 1 line crossed to *PD2* gave rise to short periods and were selected for repeats. 2 lines crossed to *TD2* and 3 lines crossed to *PD2* gave rise to long periods and were selected for repeats.

Among the 43 candidate genes (Table 1 and 2), we noticed a high proportion of genes involved or presumed to be involved in splicing (17), including five suspected or known to impact alternative splicing. Perhaps not surprisingly, several genes involved in snRNP assembly were identified in our screen. Their downregulation caused long period phenotypes. We also noticed the presence of four members of the CCR4-NOT complex, which can potentially regulate different steps of mRNA metabolism, including deadenylation, and thus mediate translational repression. Their downregulation mostly caused short period phenotypes and tended to result in high levels of arrhythmicity. *Rga* downregulation, however, resulted in a long period phenotype, suggesting multiple functions for the CCR4-NOT complex in the regulation of circadian rhythms. Interestingly, two genes implicated in mRNA decapping triggered by deadenylation, were also identified, with long periods observed when these genes were downregulated.

**Table 2:** Predicted or known functions of screen candidates.

| **Table 2: Predicted or known functions of screen candidates** |  |
| --- | --- |
| **Gene** | **Molecular Function (based on information from Flybase)** |
| Atx-1 | RNA binding |
| barc | mRNA splicing; mRNA binding; U2 snRNP binding |
| bsf | mitochondrial mRNA polyadenylation, stability, transcription, translation; polycistronic mRNA processing; mRNA 3'-UTR binding |
| CG16941/Sf3a1 | alternative mRNA splicing; RNA binding |
| CG32364/tut | translation; RNA binding |
| CG42458 | mRNA binding |
| CG4849 | mRNA splicing; translational elongation |
| CG5808 | mRNA splicing; protein peptidyl-prolyl isomerization; regulation of phosphorylation of RNA polymerase II C-terminal domain; mRNA binding |
| CG6227 | alternative mRNA splicing; ATP-dependent RNA helicase activity |
| CG7903 | mRNA binding |
| CG8273/Son | mRNA processing; mRNA splicing; RNA binding |
| CG8636/eIF3g1 | translational initiation; mRNA binding |
| CG9609 | transcription; proximal promoter sequence-specific DNA binding |
| Cnot4 | CCR4-NOT complex |
| Dcp2 | deadenylation-dependent decapping of mRNA; cytoplasmic mRNA P-body assembly; RNA binding |
| eIF1 | ribosomal small subunit binding; RNA binding; translation initiation |
| eIF3l | translational initiation |
| Hrb98DE | translation; alternative mRNA splicing; mRNA binding |
| l(1)G0007 | alternative mRNA splicing; 3'-5' RNA helicase activity |
| LSm7 | mRNA splicing; mRNA catabolic process; RNA binding |
| ncm | mRNA splicing; RNA binding |
| Nelf-A | transcription elongation; RNA binding |
| Not1 | translation; poly(A)-specific ribonuclease activity; CCR4-NOT complex |
| Not3 | translation; transcription; poly(A)-specific ribonuclease activity; CCR4-NOT complex |
| Patr-1 | cytoplasmic mRNA P-body assembly; deadenylation-dependent decapping of mRNA; RNA binding |
| Pcf11 | mRNA polyadenylation; transcription termination; mRNA binding |
| pcm | cytoplasmic mRNA P-body assembly; 5'-3' exonuclease activity |
| Psi | alternative mRNA splicing; transcription; mRNA binding |
| Rga | translation; transcription; poly(A)-specific ribonuclease activity; CCR4-NOT complex |
| RpS3 | DNA repair; translation; RNA binding; structural constituent of ribosome |
| Rrp6 | chromosome segregation; mRNA polyadenylation; nuclear RNA surveillance; 3'-5' exonuclease activity |
| sbr | mRNA export from nucleus; mRNA polyadenylation; RNA binding |
| Set1 | histone methyltransferase activity; nucleic acid binding; RRM |
| SmB | mRNA splicing; RNA binding |
| SmE | mRNA splicing; spliceosomal snRNP assembly |
| SmF | mRNA splicing; spliceosomal snRNP assembly; RNA binding |
| smg | RNA localization; translation; mRNA poly(A) tail shortening; transcription; mRNA binding |
| Smg5 | nonsense-mediated decay; ribonuclease activity |
| Smn | spliceosomal snRNP assembly; RNA binding |
| snRNP-U1-C | mRNA 5'-splice site recognition; mRNA splicing, alternative mRNA splicing |
| Spx | mRNA splicing; mRNA binding |
| Srp54k | SRP-dependent cotranslational protein targeting to membrane; 7S RNA binding |
| Zn72D | mRNA splicing; RNA binding |

### Knockdown of *Psi* shortens the period of behavioral rhythms

A promising candidate to emerge from our screen was the alternative splicing regulator *P-element somatic inhibitor*, or *Psi* (Siebel et al., 1992; Labourier et al., 2001)

The phenotype caused by *Psi* downregulation was more pronounced with *TD2* than with *PD2* (Figure 2, Table 3). This was unexpected since the sLNvs - targeted quite specifically by *PD2* - determine circadian behavior period in DD (Renn et al., 1999; Stoleru et al., 2005). This could happen because *PD2* is less efficient at downregulating *Psi* in sLNvs than *TD2*, or because the short period phenotype is not solely caused by downregulation of *Psi* in the sLNvs. To distinguish between these two possibilities, we added *Pdf-GAL80* to *TD2* to inhibit *GAL4* activity specifically in the LNvs (Stoleru et al., 2004), while allowing RNAi expression in all other circadian tissues. With this combination, we also observed a significant period shortening compared to controls, but the period shortening was not as pronounced as with *TD*2 (Figure 2D, Table 3). We therefore conclude that both the sLNvs and non-PDF cells contributes to the short period phenotype caused by *Psi* downregulation (see discussion).

**Table 3:**
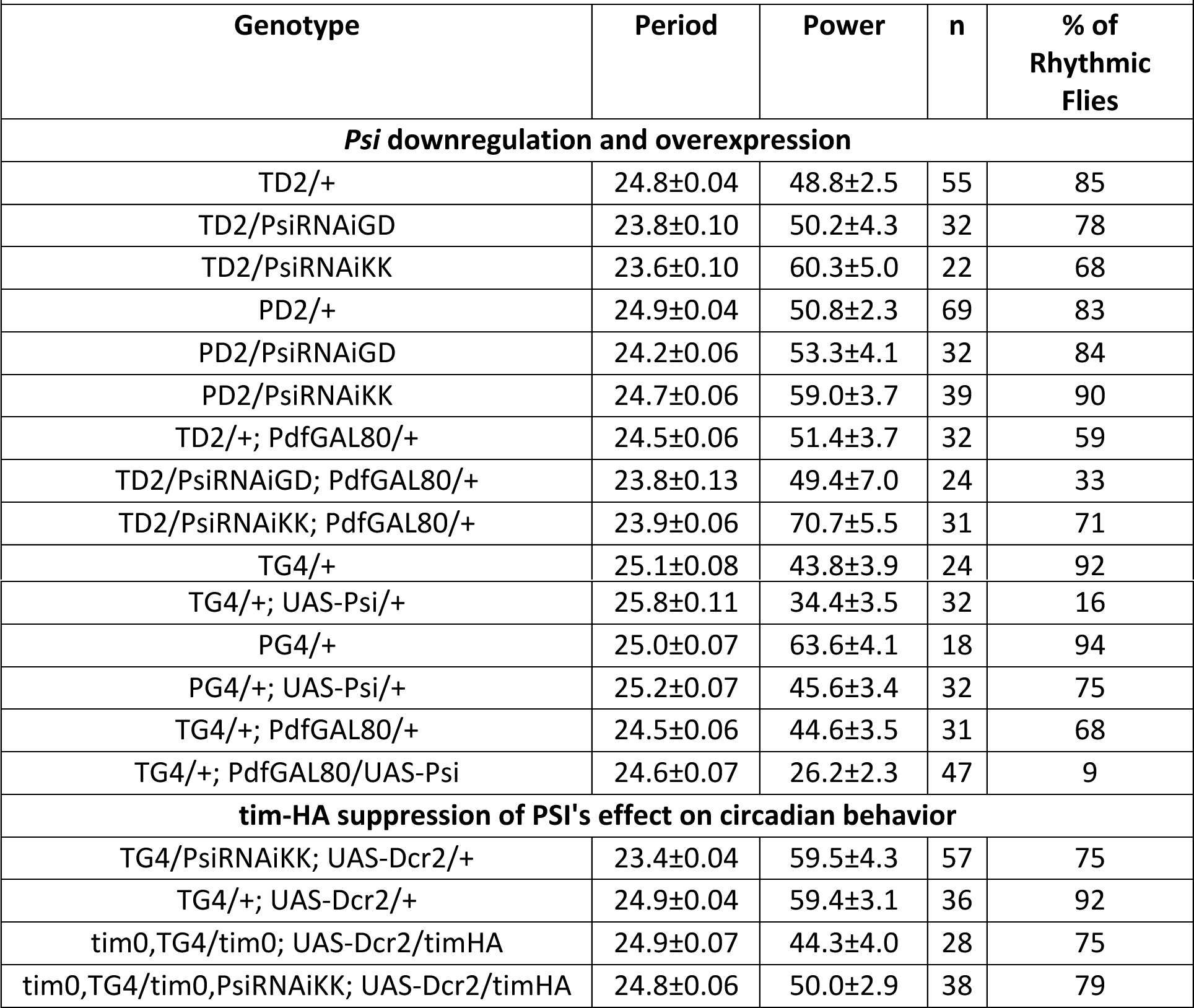
PSI affects circadian behavior.

**Figure 2.**
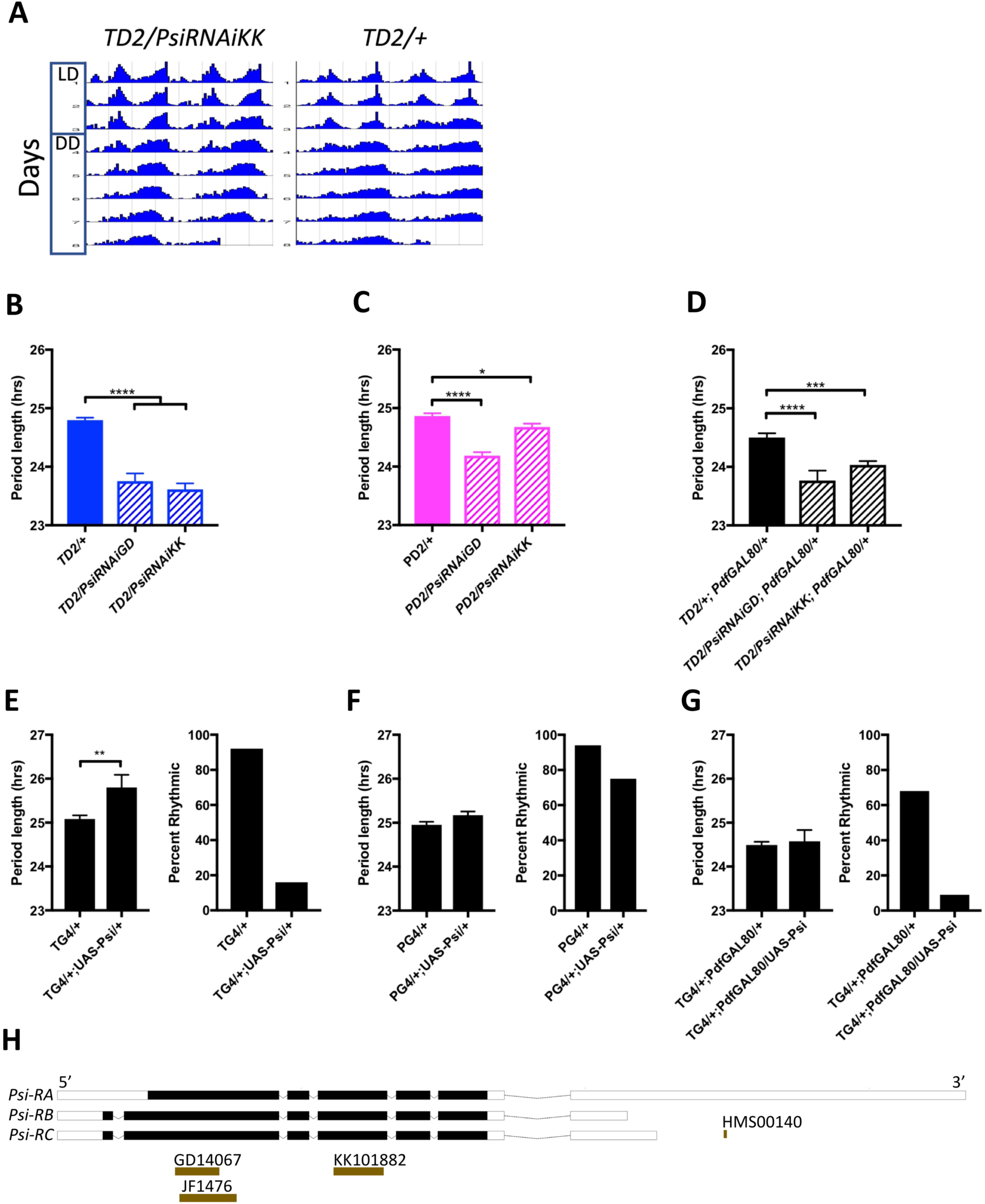
Expression level of *Psi* affects the circadian behavior period length and circadian rhythmicity. (A-D) Knockdown of *Psi* shortens the behavioral period. (A) Double-plotted actograms showing the average activities during 3 days in LD and 5 days in DD. Left panel: *TD2/PsiRNAi* (*Psi* knockdown) flies. Right panel: *TD2/+* (control) flies. Note the short period of *Psi* knockdown flies. n=8 flies/genotype. (B-D) Circadian period length (hrs) is plotted on the y axis. Genotypes are listed on the x axis. Error bars represent SEM. Solid bars are driver controls and patterned bars are *Psi* knockdown with 2 non-overlapping RNAi lines: *GD14067* (PsiRNAiGD) and *KK101882* (PsiRNAiKK). ****p<0.0001, ***p<0.001, *p<0.05, one-way analysis of variance (ANOVA) followed by Dunnett’s multiple comparison test. (B) Knockdown in all circadian tissues. (C) Knockdown in PDF+ circadian pacemaker neurons. (D) Knockdown in PDF-circadian tissues. (E-G) Overexpression of *Psi* lengthens the behavioral period and decreases rhythmicity. Left panels: Circadian period length (hrs) is plotted on the y axis. Right panels: Percent of flies that remained rhythmic in DD is plotted on the y axis. Both panels: Genotypes are listed on the x axis. Error bars represent SEM. (E) Overexpression of *Psi* in all circadian tissues lengthened the circadian period (**p<0.01, Student’s t-test) and decreased the percent of rhythmic flies. (F) Overexpression of *Psi* in PDF+ circadian pacemaker neurons caused a slight but non-significant period lengthening and a small decrease in rhythmicity. (G) Overexpression of *Psi* in PDF-circadian tissues did not affect circadian period but led to a large decrease in rhythmicity. (H) Schematic of *Psi* isoforms and position of the long and short hairpin RNAi lines used in this study.

### *Psi* overexpression disrupts circadian behavior

Since we observed that downregulating *Psi* leads to a short period, we wondered whether overexpression would have an inverse effect and lengthen the period of circadian behavior. Indeed, when we overexpressed *Psi* by driving a *UAS-Psi* transgene (Labourier et al., 2001) with the *tim-GAL4 (TG4)* driver, the period length of circadian behavior increased significantly by about 0.5 hours (Figure 2E). Interestingly, we also observed a severe decrease in the number of rhythmic flies. When we overexpressed *Psi* with *Pdf-GAL4 (PG4)*, period was not statistically different from controls, and rhythmicity was only slightly decreased (Figure 2F). Overexpression of *Psi* with the *tim-GAL4; Pdf-GAL80* combination caused a severe decrease in rhythmicity but did not lengthen period (Figure 2G). The effect of *Psi* overexpression on period is in line with the knockdown results, indicating that PSI regulates circadian behavior period through both PDF+ LNvs and non-PDF circadian neurons. However, the increase in arrhythmicity observed with *Psi* overexpression is primarily caused by non-PDF cells.

### *Psi* downregulation also shortens the period of body clocks

We wanted to further examine the effect of *Psi* knockdown on the molecular rhythms of two core clock genes: *period (per)* and *timeless (tim).* To do this, we took advantage of two *luciferase* reporter transgenes. We downregulated *Psi* with the *TD2* driver in flies expressing either a TIM-LUCIFERASE (TIM-LUC) or a PER-LUCIFERASE (BG-LUC) fusion protein under the control of the *tim* or *per* promoter, respectively. The period of LUC activity was significantly shortened by about 1-1.5 hours compared to controls when *Psi* was downregulated in TIM-LUC flies (Figure 3A and 3B). This is consistent with the behavioral period shortening we observed in *TD2>PsiRNAi* flies. Knockdown of *Psi* in BG-LUC flies resulted in a similar trend, although differences did not reach statistical significance (Figure 3C and 3D). Since the LUCIFERASE signal in these flies is dominated by light from the abdomen, this indicates that *Psi* knockdown, shortens the period of circadian clocks in peripheral tissues as well as in the brain neural network that controls circadian behavior.

**Figure 3.**
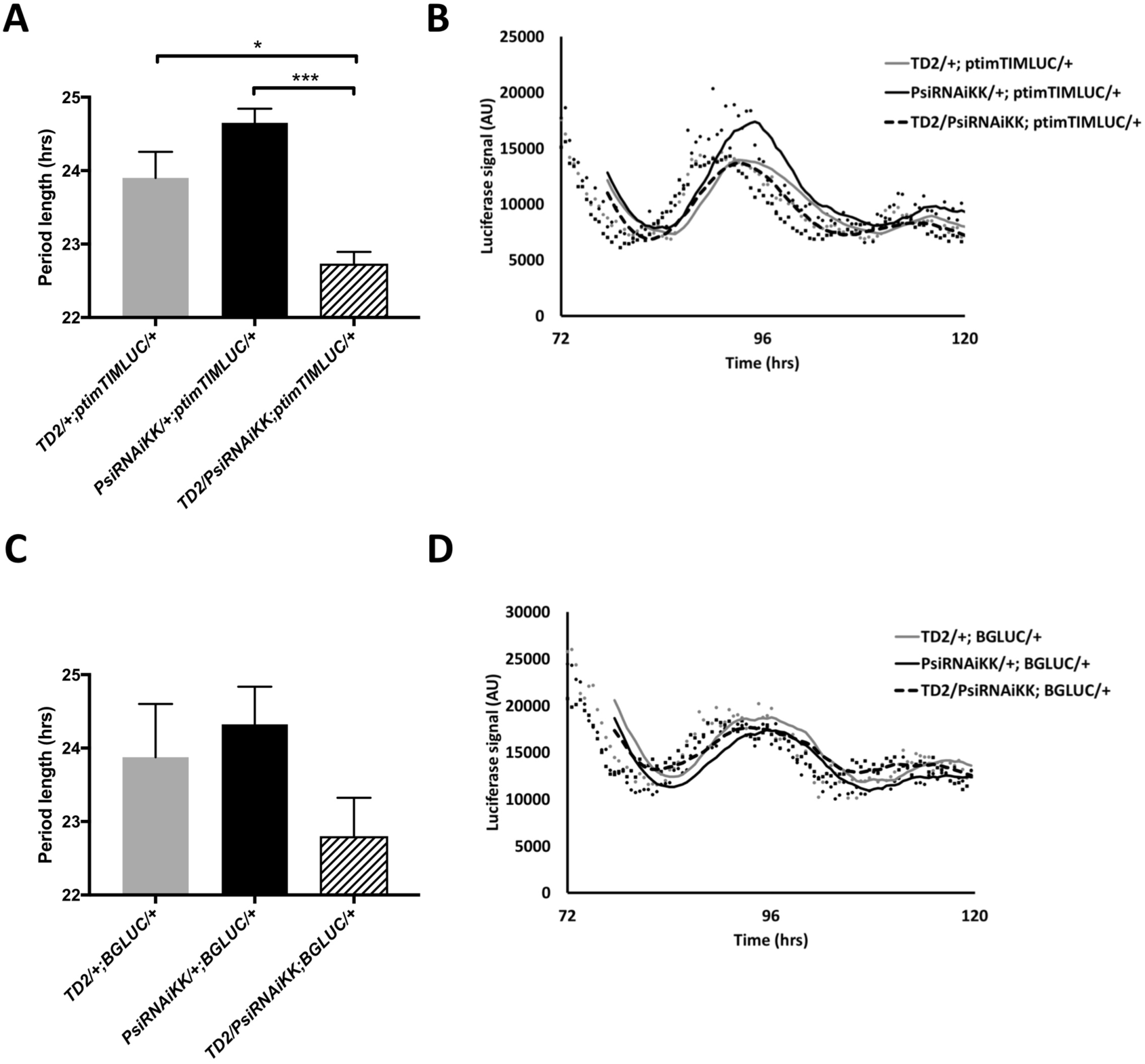
Knockdown of Psi shortens circadian period of PER and TIM rhythms in peripheral tissues. (A) Period length (hrs) of light output generated from luciferase rhythms of *ptim*-TIM-LUC in whole flies. 12-24 wells/run, 3 flies/well. N=6 runs. *p<0.05, ***p<0.001, one-way analysis of variance (ANOVA) followed by Tukey’s multiple comparison test. Error bars represent SEM. Gray bar: driver control. Black bar: RNAi control. Patterned bar: *Psi* knockdown. (B) Representative traces from (A) Markers are raw data and lines are 6-hour moving averages. Gray marker (circle) and gray line: driver control. Black marker (circle) and black line: RNAi control. Black marker (square) and dashed line: *Psi* knockdown. Luciferase signal (arbitrary units, AU) on the y axis and time (hrs) from start of experiment on the x axis. 72 hrs = start of DD. (C) Period length (hrs) of average light output generated from luciferase rhythms of BG-LUC in whole flies. 12-24 wells/run, 3 flies/well. N=4 runs. Error bars represent SEM. Gray bar: driver control. Black bar: RNAi control. Patterned bar: *Psi* knockdown. (D) Representative traces from (C) Markers are raw data and lines are 6-hour moving averages. Gray marker (circle) and gray line: driver control. Black marker (circle) and black line: RNAi control. Black marker (square) and dashed line: *Psi* knockdown. Luciferase signal (arbitrary units, AU) on the y axis and time (hrs) from start of experiment on the x axis. 72 hrs = start of DD.

### Alternative splicing of two clock genes, *cwo* and *tim*, is altered in *Psi* knockdown flies

PSI has been best characterized for its role in alternative splicing of the *P element transposase* gene in somatic cells (Siebel et al., 1992; Labourier et al., 2001). However, it was recently reported that PSI has a wider role in alternative splicing (Wang et al., 2016). Wang *et al* reported an RNA-seq dataset of alternative splicing changes that occur when a lethal *Psi-*null allele is rescued with a copy of *Psi* in which the AB domain has been deleted (PSIΔAB). This domain is required for the interaction of PSI with the U1 snRNP, which is necessary for PSI to mediate alternative splicing of *P element* transposase (Labourier et al., 2002). Interestingly, Wang *et al* found that *Psi* affects alternative splicing of genes involved in complex behaviors such as learning, memory and courtship (Wang et al., 2016). Intriguingly, we found four core clock genes listed in this dataset: *tim*, *cwo*, *sgg* and *Pdp1*. We decided to focus on *cwo* and *tim*, since only one specific splicing isoform of *Pdp1* is involved in the regulation circadian rhythm, (*Pdp1e*) (Zheng et al., 2009), and since the *sgg* gene produces a very complex set of alternative transcripts. After three days of LD entrainment, we collected RNA samples at four time points on the first day of DD and determined the relative expression of multiple isoforms of *cwo* and *tim* in *Psi* knockdown heads compared to driver and RNAi controls.

CWO is a bHLH transcriptional factor and is part of an interlocked feedback loop that reinforces the main loop by competing with CLK/CYC for E-box binding (Kadener et al., 2007; Lim et al., 2007; Matsumoto et al., 2007; Richier et al., 2008). There are three mRNA isoforms of *cwo* predicted in Flybase. Of the three, only *cwoRA* encodes a full-length CWO protein. Exon 2 is skipped in *cwoRB*, and in *cwoRC* there is an alternative 3’ splice site in the first intron that lengthens exon 2. Both *cwo RB and RC* have an alternative 3’ splice site in exon 3, and translation begins from a downstream start codon in that exon (flybase). The predicted start codon in both *cwoRB* and *cwoRC* would produce an N-terminal truncation of the protein that would be missing the basic region of the bHLH domain and should thus not be able to bind DNA. The *cwoRB* and cwo*RC* isoforms may thus act as endogenous dominant negatives.

We found that the levels of the *cwoRA* and *cwoRB* isoforms were significantly reduced compared to both controls at CT 9 (Figure 4A and B). Conversely, the *cwoRC* isoform expression was significantly increased at CT 15 (Figure 4C). The overall expression of all *cwo* mRNAs in *Psi* knockdown fly heads was significantly reduced at both CT 9 and CT 15, indicating that the RC isoform’s contribution to total *cwo* mRNA levels is quite modest (Figure 4D).

**Figure 4.**
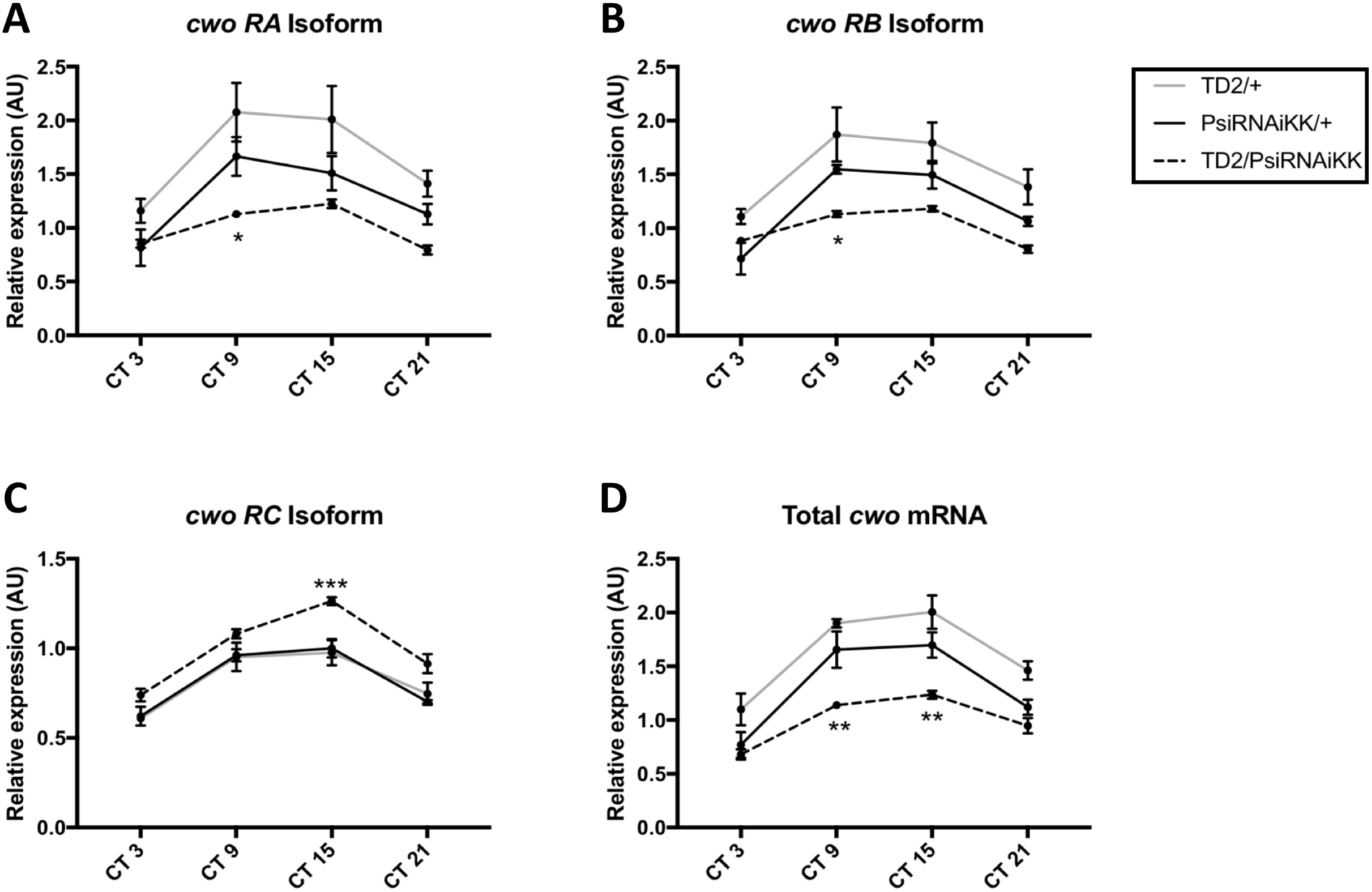
Knockdown of Psi affects the balance of *cwo* isoform expression. (A-C) Relative expression of *cwo* mRNA isoforms (normalized to the average of all *Psi* knockdown time points) in heads on the y axis measured by qPCR. (D) Relative expression of total *cwo* mRNA on the y axis. (A-D) Circadian time (CT) on the x axis. Error bars represent SEM. Gray line: driver control. Black line: RNAi control. Dashed line: *Psi* knockdown. Driver control, N=3. RNAi control, N=4. *Psi* knockdown, N=6. Both driver and RNAi control relative to *Psi* knockdown, two-way analysis of variance (ANOVA) followed by Tukey’s multiple comparison test: *p<0.05, **p<0.01, ***p<0.001.

We then analyzed alternative splicing of *tim* in *Psi* knockdown heads compared to controls. Specifically, we looked at the expression of three temperature-sensitive intron inclusion events in *tim* that all theoretically lead to C-terminal truncations of the protein. The *tim-cold* isoform has been previously described. This isoform, which is dominant at low temperature (18°C), arises when the last intron is included (Boothroyd et al., 2007). We found that *tim-cold* is elevated in *Psi* knockdown heads at peak levels under 25°C conditions (CT15, Figure 5B). Similarly, we found that another intron inclusion event, *tim-sc* which is also elevated at 18°C and is present in the *timRN* and *RO* isoforms (see accompanying paper by Evantal et al.), is significantly increased at 25°C in *Psi* knockdown heads at CT15 (Figure 5A). Thus, interestingly, two intron inclusion events that are upregulated by cold temperature are also both upregulated in *Psi* knockdown heads at 25°C. In contrast, we found that an intron included in the *timRM* and *RS* isoforms (*tim-M*) and increased at high temperature (29°C, see accompanying paper by Evantal et al.) is significantly decreased at CT 9, 15 and 21 in *Psi* knockdown heads at 25°C (Figure. 5C). It should be noted that the *tim-sc* intron is only partially retained, because a cleavage and poly-adenylation signal is located within this intron, thus resulting in a much shorter mature transcript (see accompanying paper by Evantal et al.). Based on PSI function, the most parsimonious explanation is that PSI reduces production of *tim-sc* by promoting splicing of the relevant intron. However, we cannot entirely exclude that PSI regulates the probability of the RNA polymerase to undergo premature transcription termination soon after passing the poly-adenylation signal.

**Figure 5.**
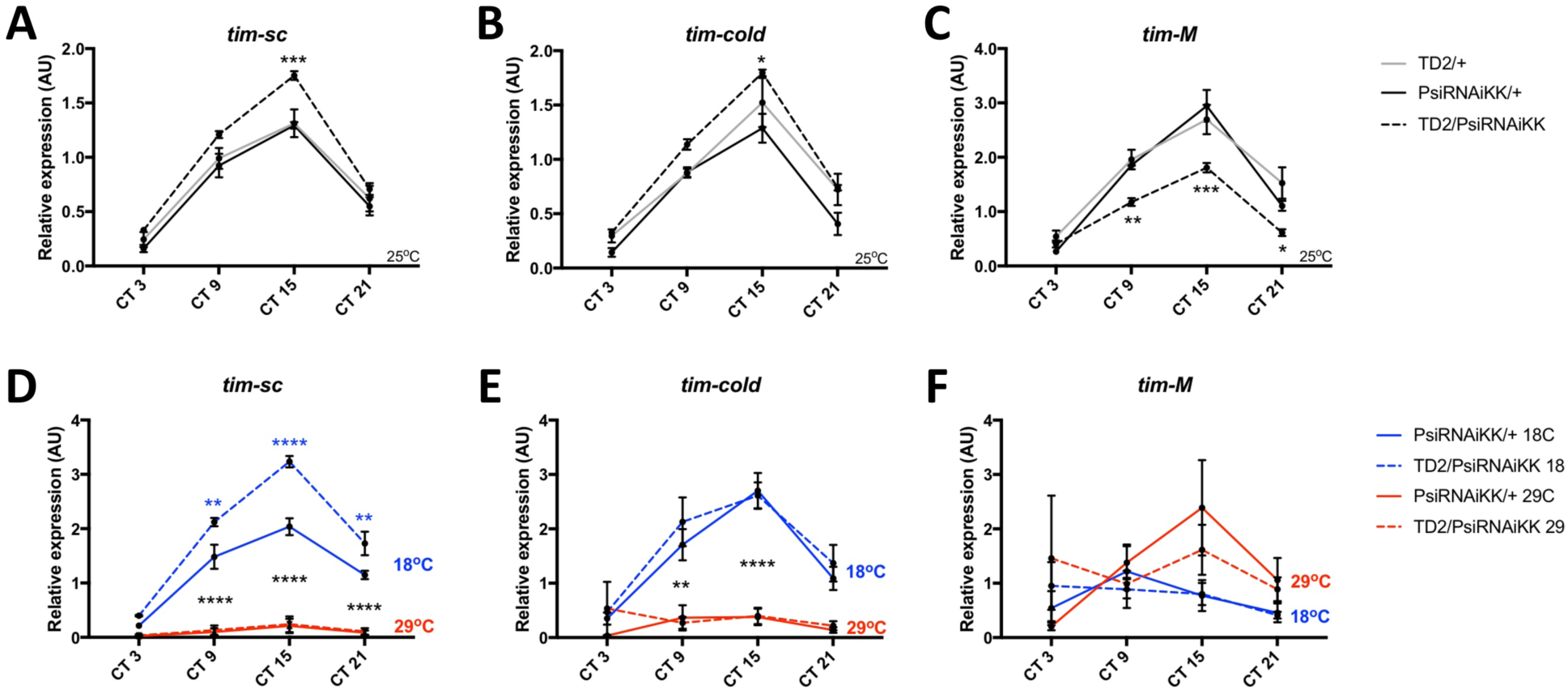
Knockdown of Psi increases the expression of cold induced *tim* isoforms and decreases the expression of a warm induced *tim* isoform. (A-C) Relative expression of *tim* mRNA isoforms at 25°C (normalized to the average of all *Psi* knockdown time points) in heads on the y axis measured by qPCR. Circadian time (CT) on the x axis. Error bars represent SEM. Gray line: driver control. Black line: RNAi control. Dashed line: *Psi* knockdown. Controls, N=3. *Psi* knockdown, N=5. Both driver and RNAi control relative to *Psi* knockdown, two-way analysis of variance (ANOVA) followed by Tukey’s multiple comparison test: *p<0.05, **p<0.01, ***p<0.001, ****p<0.0001. (D-F) Relative expression of *tim* mRNA isoforms at 18°C and 29°C Solid line: RNAi control. Dashed line: *Psi* RNAi knockdown. Blue indicates flies were transferred to 18°C at CT0 (start of subjective day) on the first day of DD. Red indicates flies were transferred to 29°C. N=3. 18°C samples compared to 29°C samples, *p<0.05, **p<0.01, ***p<0.001, ****p<0.0001, two-way analysis of variance (ANOVA) followed by Tukey’s multiple comparison test. (D) Blue asterisks refer to RNAi control compared to *Psi* knockdown.

Collectively, these results indicate that PSI shifts the balance toward a warm temperature *tim* RNA isoform profile at an intermediate temperature (25°C). This could be achieved either by altering the temperature sensitivity of *tim* introns, or by promoting a “warm temperature splicing pattern” independently of temperature. We therefore also measured *tim* splicing isoforms at 18 and 29°C (Figure 5D-F). We entrained flies for 3 days in LD at 25°C to maintain similar levels of GAL4 expression and thus of *Psi* knockdown (the GAL4/UAS system’s activity increases with temperature (Duffy, 2002). We then shifted them to either 18°C or 29°C at CT 0 on the first day of DD and collected samples at CT 3, 9, 15 and 21. We found that both the *tim-cold* intron and the *tim-sc* introns were elevated at 18°C in both *Psi* knockdown heads and controls (Figure 5D and E). Thus, *Psi* knockdown does not block the temperature sensitivity of these introns. *tim-M* levels were unexpectedly variable in DD, particularly in the *Psi* knockdown flies, perhaps because of the temperature change. Nevertheless, we observed a trend for the *tim-M* intron retention to be elevated at 29°C (Figure 5F), further supporting our conclusion that *Psi* knockdown does not affect the temperature sensitivity of *tim* splicing, but rather, the balance of isoform expression at all temperatures.

As expected from these results, *Psi* downregulation did not affect the ability of flies to adjust the phase of their evening and morning peak to changes in temperature (Figure S2). We also tested whether *Psi* knockdown flies responded normally to short light pulses, since TIM is the target of the circadian photoreceptor CRY (Emery et al., 1998; Stanewsky et al., 1998; Lin et al., 2001; Busza et al., 2004; Koh et al., 2006). These flies could both delay or advance the phase of their circadian behavior in response to early or late-night light pulses, respectively (Figure S3). We noticed however a possible slight shift of the whole Phase Response Curve toward earlier times. This would be expected since the pace of the circadian clock is accelerated.

### *tim* splicing is required for PSI’s regulation of circadian period

Because *tim* is a key element of the circadian transcriptional feedback loop, we wondered whether *Psi* might be regulating the speed of the clock through its effects on *tim* splicing. We therefore rescued the amoprhic *tim* allele (*tim^0^)* with a *tim* transgene that lacks the known temperature sensitive alternatively spliced introns as well as most other introns (*tim-HA*) (Figure 6A) (Rutila et al., 1998). Importantly, the *tim^0^* mutation is a frame-shifting deletion located upstream of the temperature-sensitive alternative splicing events (Myers et al., 1995), and would thus truncate any TIM protein produced from the splice variants we studied. Strikingly, we found that knockdown of *Psi* in *tim-HA* rescued *tim^0^* flies had no impact on the period of circadian behavior (Figure 6B, Table 3). This indicates that PSI controls circadian period through *tim* splicing.

**Figure 6.**
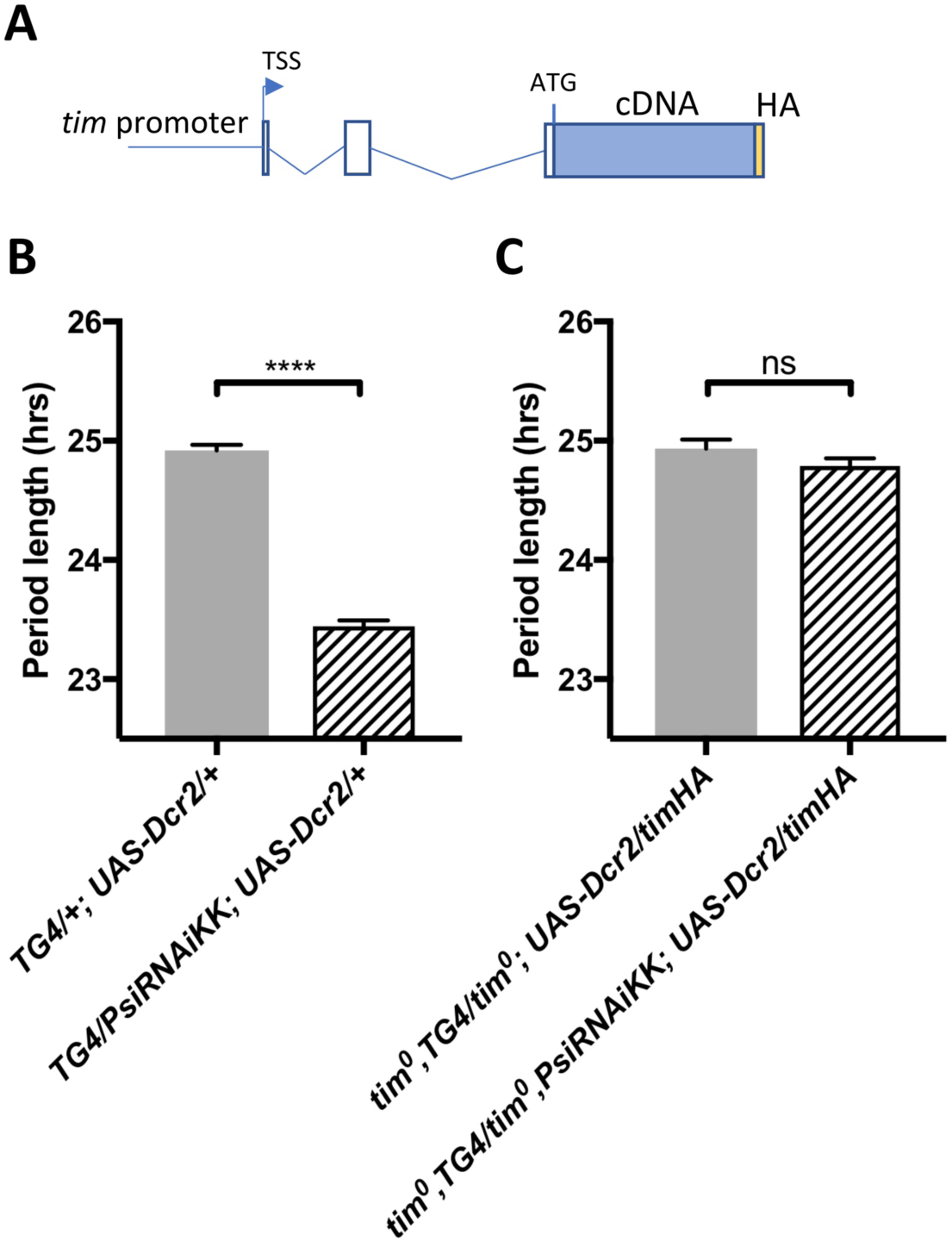
The short period effect of Psi knockdown is dependent on tim introns. (A) Schematic of *timHA* transgene. The *tim* promoter is fused upstream of the transcription start site (TSS). Two introns remain in the 5’UTR, upstream of the start codon; however, they are not, to our knowledge, temperature sensitive. A C-terminal HA tag is fused to full length *tim* cDNA, which lacks any of the introns that are known to be retained at high or low temperatures. (B) Knockdown of *Psi* with *TD2* causes period shortening. ****p<0.0001, Student’s t-test. (C) Period shortening in response to *Psi* knockdown with *TD2* is abolished in *tim^0^, ptim-timHA* flies that can only produce the full length *tim* isoform. ns, p=0.1531, Student’s t-test. (B-C) Circadian period length (hrs) is plotted on the y axis. Genotypes are listed on the x axis. Error bars represent SEM.

## Discussion

Our results identify a novel post-transcriptional regulator of the circadian clock: *Psi*. *Psi* is required for the proper pace of both brain and body clock. When *Psi* is downregulated, the circadian pacemaker speeds up, and this appears to be predominantly caused by an abnormal *tim* splicing pattern. Indeed, the circadian period of flies that can only produce functional TIM protein from a transgene missing most introns is insensitive to *Psi* downregulation. We note however that *cwo*’s splicing pattern is also affected by *Psi* downregulation, and we did not study *sgg* splicing pattern, although it might also be controlled by PSI (Wang et al., 2016). We therefore cannot exclude a small contribution of non-*tim* splicing events to period changes, or that in specific tissues these other splicing events play a greater role than in the brain.

Interestingly, PSI downregulation results in an increase in intron inclusion events that are favored under cold conditions, while an intron inclusion event favored under warm conditions is decreased. However, the ability of *tim* splicing to respond to temperature changes is not altered when *Psi* is downregulated (Fig 5D-F). This could imply that an as yet unknown factor specifically promotes or represses *tim* splicing events in a temperature-dependent manner. Another possibility is that the strength of splice sites or *tim*’s pre-mRNA structure impacts splicing efficiency in a temperature–dependent manner. For example, suboptimal *per* splicing signals explain the higher efficiency of *per*’s most 3’ splicing event at cold temperature (Low et al., 2008).

How would the patterns of *tim* splicing affect the pace of the circadian clock? In all splicing events that we studied, the result of intron retention is a truncated TIM protein. It is therefore possible that the balance of full length and truncated TIM proteins determine circadian period. Consistent with this idea, overexpression of the shorter cold-favored TIM isoform shortens period (see accompany paper by Kadener et al). Strikingly, *Psi* downregulation increases this isoform’s levels and also results in a short phenotype.

Other splicing factors have been shown to be involved in the control of circadian rhythms in *Drosophila*. SRm160 contributes to the amplitude of circadian rhythms by promoting *per* expression (Beckwith et al., 2017), while B52/SMp55 and PRMT5 regulate *per’s* most 3’ splicing, which is temperature sensitive (Sanchez et al., 2010; Zhang et al., 2018). Loss of PRMT5 results in essentially arrhythmic behavior (Sanchez et al., 2010), but this is unlikely to be explained by its effect on *per’s* thermosensitive splicing. B52/SMp55 show a reduced siesta, which is controlled by the same *per* splicing (Zhang et al., 2018). With the identification of *Psi*, we uncover a key regulator of *tim* alternative splicing pattern and show that this pattern determines circadian period length, while *per* alternative splicing regulates the timing and amplitude of the daytime siesta. Interestingly, a very recent study identified PRP4 kinase and other members of tri-snRNP complexes as regulators of circadian rhythms (Shakhmantsir et al., 2018). Downregulation of PRP4 caused excessive retention of *tim-M* (called *tim-tiny* in that study).

An unexpected finding is the role played by both PDF neurons and other circadian neurons in the short period phenotype observed with circadian locomotor rhythms when we knocked-down PSI. Indeed, it is quite clear from multiple studies that under constant darkness, the PDF-positive sLNvs dictate the pace of circadian behavior (Stoleru et al., 2005; Yao and Shafer, 2014). Why, in the case of Psi downregulation, do PDF negative neurons also play a role in period determination? The explanation might be that *Psi* alters the hierarchy between circadian neurons, promoting the role of PDF negative neurons. This could be achieved by weakening PDF/PDFR signaling, for example.

While we focused our work on PSI, several other interesting candidates were identified in our screen (Table 1 and 2). We note the presence of a large number of splicing factors. This adds to the emerging notion that alternative splicing plays a critical role in the control of circadian rhythms. We have already mentioned above several *per* splicing regulators that can impact circadian behavior. In addition, a recent study demonstrate that specific classes of circadian neurons express specific alternative splicing variants, and that rhythmic alternative splicing is widespread in these neurons (Wang et al., 2018). Interestingly, in this study, the splicing regulator *barc*, which was identified in our screen and which has been shown to causes intron retention in specific mRNAs (Abramczuk et al., 2017), was found to be rhythmically expressed in LNds. Moreover, in mammals alternative splicing appears to be very sensitive to temperature, and could explain how body temperature rhythms synchronize peripheral clocks (Preußner et al., 2017). Another intriguing candidate is *cg42458*, which was found to be enriched in circadian neurons (LNvs and Dorsal Neurons 1) (Wang et al., 2018). In addition to emphasizing the role of splicing, our screen suggests that regulation of polyA tail length is important for circadian rhythmicity, since we identified several members of the CCR4-NOT complex and deadenylation-dependent decapping enzymes. Future work will be required to determine whether these factors directly target mRNAs encoding for core clock components, or whether their effect on circadian period is indirect. It should be noted that while our screen targeted 364 proteins binding or associated with RNA, it did not include all of them. For example, LSM12, which was recently shown to be a part of the ATXN2/TYF complex (Lee et al., 2017), was not included in our screen because it had not been annotated as a potential RAP when we initiated our screen.

In summary, our work provides an important resource for identifying RNA associated proteins regulating circadian rhythms in *Drosophila*. It identifies PSI is an important regulator of circadian period, and points at additional candidates and processes that determine the periodicity of circadian rhythms.

## Materials and Methods

### Fly stocks

Flies were raised on a standard cornmeal/agar medium at 25°C under a 12hr:12hr light:dark (LD) cycle. The following *Drosophila* strains were used: *w^1118^ -- w; Tim-GAL4, UAS-dicer2/CyO (TD2)* (Dubriulle et al., 2009) *-- y w; Pdf-GAL4, UAS-dicer2/CyO (PD2)* (Dubriulle et al., 2009) *-- y w; Tim-GAL4/CyO (TG4)* (Kaneko et al., 2000) *-- y w; Pdf-GAL4 (PG4)* (Renn et al., 1999) *---- w;; UAS-dcr2* (Dietzl et al., 2007) *---- y w;; timHA* (Rutila et al., 1998). The following combinations were generated for this study: *w; tim-GAL4/CyO; UAS-dicer2/TM6B -- tim^0^,TG4/CyO; UAD-Dcr2/TM6B -- tim^0^, PsiRNAiKK/CyO; timHA/TM6B. TD2, ptim-TIM-LUC* and *TD2, BG-LUC* transgenic flies expressing a *tim-luciferase* and *per-luciferase* fusion gene respectively, combined with the TD2 driver, were used for luciferase experiments. The TIM-LUC fusion is under the control of the *tim* promoter (ca. 5kb) and 1^st^ intron (Lamba et al., 2018), BG-LUC contains per genomic DNA encoding the N-terminal two-thirds of PER and is under the control of the *per* promoter (Stanewsky et al., 1997). RNAi lines (names beginning with JF, GL, GLV, HM or HMS) were generated by the Transgenic RNAi Project at Harvard Medical School (Boston, MA) and obtained from the Bloomington Drosophila Stock Center (Indiana University, USA). RNAi lines (names beginning with GD or KK) were obtained from the Vienna Drosophila Stock Center. *UAS-Psi* flies were kindly provided by D. Rio (Labourier et al., 2001).

### Behavioral monitoring and analysis

The locomotor activity of individual male flies (2-5 days old at start of experiment) was monitored in Trikinetics Activity Monitors (Waltham, MA). Flies were entrained to a 12:12 LD cycle for 3-4 days at 25°C using I-36LL Percival incubators (Percival Scientific, Perry IA). After entrainment, flies were released into DD for five days. Rhythmicity and period length were analyzed using the FaasX software (courtesy of F. Rouyer, Centre National de la Recherche Scientifique, Gif-sur-Yvette, France) (Grima et al., 2002). Rhythmicity was defined by the criteria – power ≥20, width ≥1.5 using the χ2 periodogram analysis. For phase-shifting experiments, groups of 16 flies per genotype were entrained to a 12:12 LD cycle for 5-6 days at 25°C exposed to a 5-minute pulse of white fluorescent light (1500 lux) at different time points on the last night of the LD cycle. A separate control group of flies was not light-pulsed. Following the light pulse, flies were released in DD for six days. To determine the amplitude of photic phase shifts, data analysis was done in MS Excel using activity data from all flies, including those that were arrhythmic according to periodogram analysis. Activity was averaged within each group, plotted in Excel, and then fitted with a 4-hour moving average. A genotype-blind observer quantified the phase shifts. The peak of activity was found to be the most reliable phase marker for all genotypes. Phase shifts were calculated by subtracting the average peak phase of the light-pulsed group from the average peak phase of non-light pulsed group of flies.

### Statistical analysis

For the statistical analysis of behavioral and luciferase period length, Student’s t-test was used to compare means between two groups, and one-way analysis of variance (ANOVA), coupled to post hoc tests, was used for multiple comparisons. Tukey’s post hoc test was used when comparing three or more genotypes and Dunnett’s post hoc test was used when comparing two experimental genotypes to one control. For the statistical analysis of qPCR and the behavioral phase-shifting experiments, two-way analysis of variance (ANOVA), coupled to Tukey’s post hoc test, was used for multiple comparisons.

### Luciferase experiments

The luciferase activity of whole male flies on Luciferin (Gold-biotech) containing agar/sucrose medium (170μl volume, 1% agar, 2% sucrose, 25mM luciferin), was monitored in Berthold LB960 plate reader (Berthold technologies, USE) in l-36LL Percival incubators with 90% humidity (Percival Scientific, Perry IA). Three flies per well were covered with needle-poked Pattern Adhesive PTFE Sealing Film (Analytical sales & services 961801). The distance between the agar and film was such that the flies were not able to move vertically. Period length was determined from light measurements taken during the first two days of DD. The analysis was limited to this window because TIM-LUC and BG-LUC oscillations severely dampened after the second day of DD. Period was estimated by an exponential dampened cosinor fit using the least squares method in MS Excel (Solver function).

### Real-time quantitative PCR

Total RNA from about 30 or 60 fly heads collected at CT 3, CT9, CT15 and CT21 on the first day of DD were prepared using Trizol (Invitrogen) and Zymo Research Direct-zol RNA MiniPrep kit (R2050) following manufacturer’s instructions. 1μg of total RNA was reverse transcribed using Bio-RAD iSCRIPT cDNA synthesis kit (1708891) following manufacturer’s instructions. Real-time PCR analysis was performed using Bio-RAD iTaq Universal SYBR Green Supermix (1725121) in a Bio-RAD C1000 Touch Thermal Cycler instrument. A standard curve was generated for each primer pair, using RNA extracted from wild-type fly heads, to verify amplification efficiency. Data were normalized to *rp32* (Dubriulle et al., 2009) using the 2^−ΔΔCt^ method. Primers used: *psi-*forward GGTGCCTTGAATGGGTGAT; *psi*-reverse CGATTTATCCGGGTCCTCG; *timRM/RS-*forward TGGGAATCTCGCCCGAAAC;

*timRM/RS-*reverse AGAAGGAGGAGAAGGAGAGAGG; *timRN/RO-*forward ACTGTGCGATGACTGGTCTG;

*timRN/RO-*reverse TGCTTCAAGGAAATCTTCTG; *tim^cold^-*forward CCTCCATGAAGTCCTCGTTCG;

*tim^cold^-*reverse ATTGAGCTGGGACACCAGG; *cwo*-foward TTCCGCTGTCCACCAACTC; *cwo*-reverse CGATTGCTTTGCTTTACCAGCTC; *cwoRA*-forward TCAAGTATGAGAGCGAAGCAGC; *cwoRA*-reverse TGTCTTATTACGTCTTCCGGTGG;

*cwoRB*-forward GTATGAGAGCAAGATCCACTTTCC; *cwoRB*-reverse GATGATCTCCGTCTTCTCGATAC; *cwoRC*-forward GTATGAGAGCCAAGCGACCAC; *cwoRC*-reverse CCAAATCCATCTGTCTGCCTC.

## Acknowledgments

We are particularly grateful to Vincent van der Vinne for his help with the analysis of the luciferase recording. We also express our gratitude to Monika Chitre for help with qPCR, and Diana Bilodeau-Wentworth, Dianne Szydlik, Chunyan Yuan and Vinh Phan for technical assistance. We also thank Dr. Donald Rio as well as the Bloomington and Vienna Drosophila Resource Centers for fly stocks. This work was supported by a MIRA award from the National Institute of General Medicine Sciences (1R35GM118087) to PE, and an ERC Starting Grant to SK.

---

**Figure S1.**
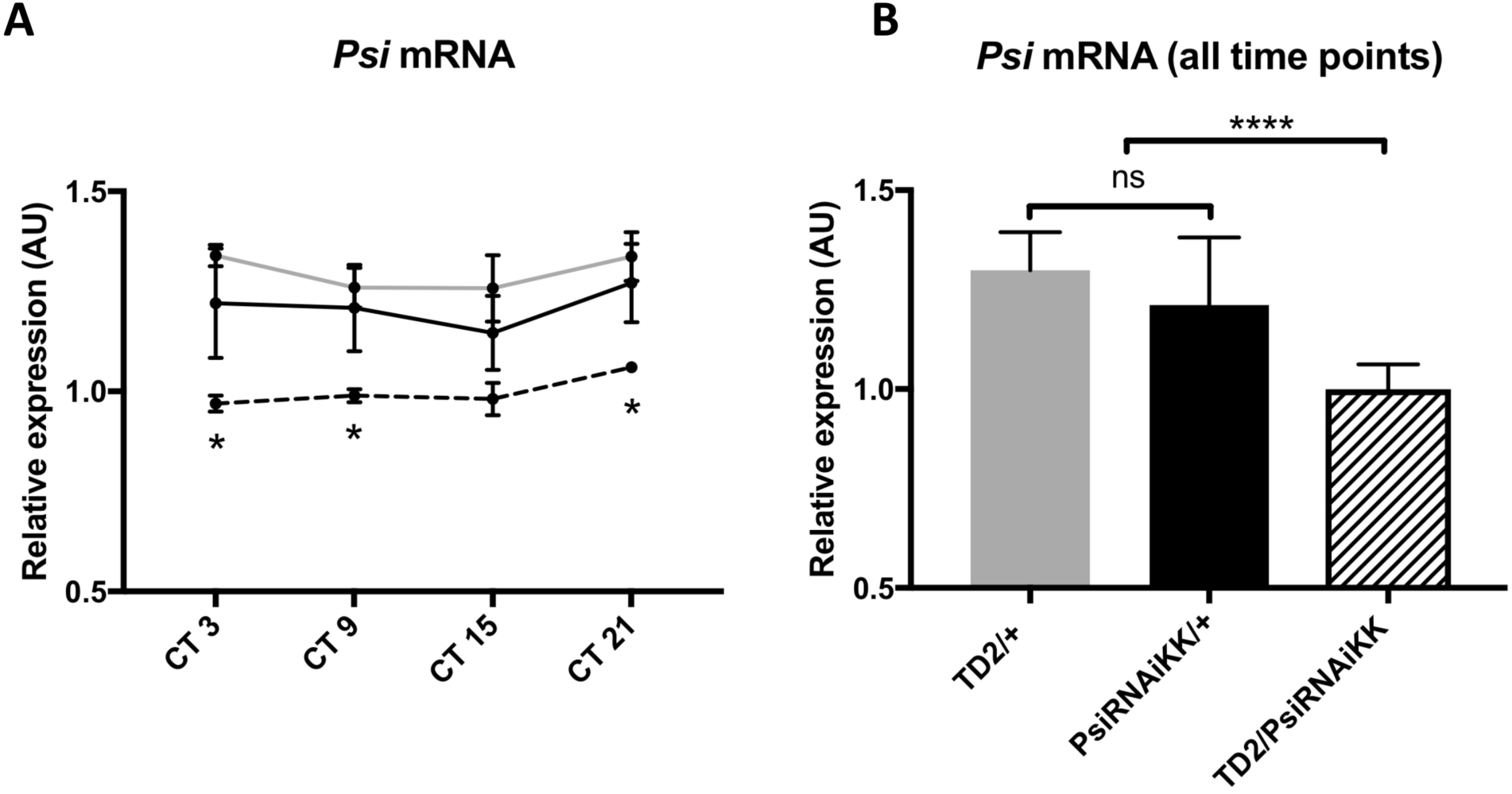
*Psi* mRNA expression does not cycle and its level is reduced in heads of *Psi* knockdown flies. *(A) Psi* mRNA expression does not cycle in DD. Relative expression of *Psi* mRNA (normalized to the average of all *Psi* knockdown time points) in heads on the y axis measured by qPCR. Circadian time (CT) on the x axis. Error bars represent SEM. Gray line: driver control. Black line: RNAi control. Dashed line: *Psi* knockdown. Controls, N=3. *Psi* knockdown, N=5. Both driver and RNAi control relative to *Psi* knockdown, two-way analysis of variance (ANOVA) followed by Tukey’s multiple comparison test: *p<0.05. (B) Knockdown of *Psi* with RNAiKK causes a significant reduction in *Psi* mRNA levels relative to both driver and RNAi controls. Since no cycling of *Psi* was observed, all time points were pooled to increase statistical strength. Relative expression of *Psi* mRNA (normalized to the average of all *Psi* knockdown time points) in heads on the y axis measured by qPCR. Genotypes are on the x axis. Error bars represent SEM. Gray bar: driver control. Black bar: RNAi control. Patterned bar: *Psi* knockdown. Both driver and RNAi control relative to *Psi* knockdown, one-way analysis of variance (ANOVA) followed by Tukey’s multiple comparison test: ****p<0.0001.

**Figure S2.**
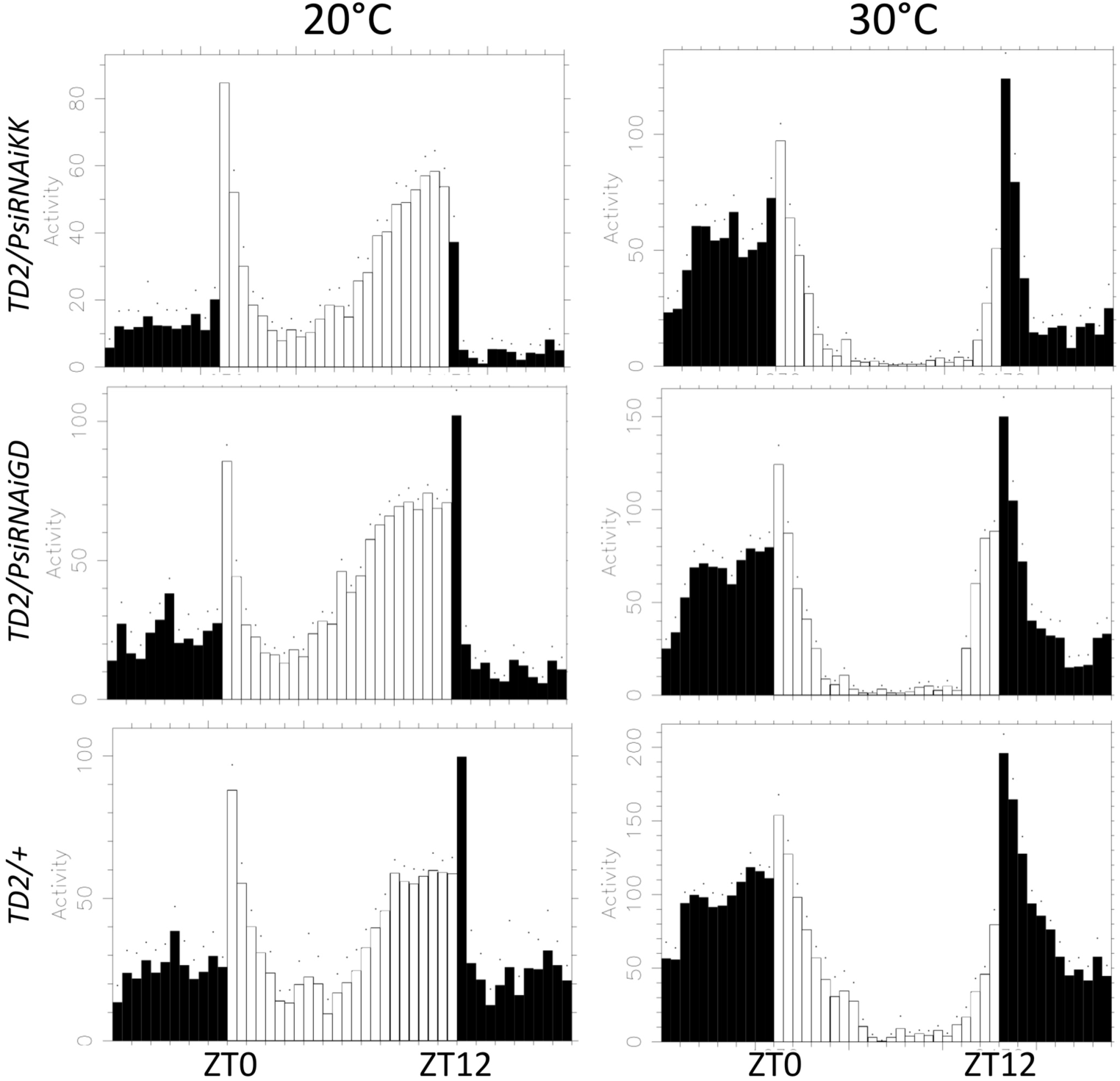
*Psi* knockdown flies have normal behavioral adaptation to temperature. Eductions showing the average activity of flies during 2 days of LD (days 2-3). Left panels: flies were entrained at 20°C. Right panels: flies were entrained at 30°C. Top panels: *TD2/PsiRNAiKK* (*Psi* knockdown) flies. Middle panels: *TD2/PsiRNAiGD* (*Psi* knockdown) flies. Bottom panels: *TD2/+* (control) flies. Note that, similar to the *TD2/+* control, *Psi* knockdown flies advance the phase of their evening activity at 20°C and delay the phase of their evening activity at 30°C. *Psi* knockdown flies also show reduced morning activity and increased evening activity at 20°C, and increased morning activity and decreased evening activity at 30°C, similar to the *TD2/+* control.

**Figure S3.**
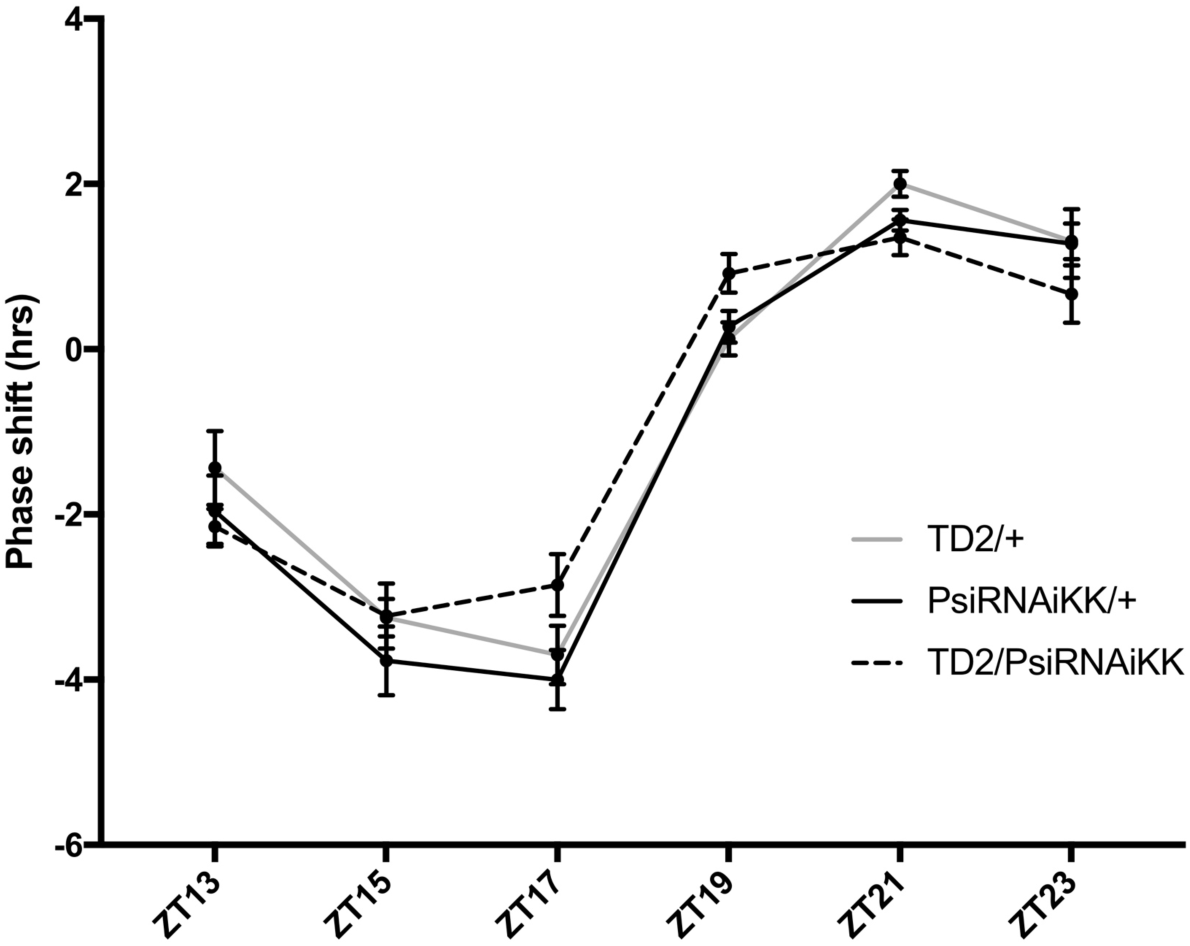
*Psi* knockdown flies have a normal photic phase response. Behavioral phase response curve to brief 5-minute 1500 lux light pulses. Behavioral phase shifts are on the y-axis. The time of the light pulse administration is on the x-axis. N = 4 for all time points except ZT23 where N = 3. For each genotype, 16 flies per timepoint were tested in each run. No significant differences were detected between *Psi* knockdown flies and controls, two-way analysis of variance (ANOVA) followed by Tukey’s multiple comparison test. Note that the phase of the *Psi* knockdown curve is slightly shifted to the left, which probably reflects the short period of *Psi* knockdown flies.

